# Experimental and Biophysical Modeling of Transcription and Translation Dynamics in Bacterial- and Mammalian-based Cell-Free Expression Systems

**DOI:** 10.1101/2021.11.12.468406

**Authors:** Yuwen Zhao, Shue Wang

**Affiliations:** Department of Chemistry, Chemical and Biomedical Engineering, Tagliatela College of Engineering, University of New Haven, West Haven, CT, 06516, USA; Department of Biomedical Engineering, Lehigh University, Bethlehem, PA, 18015, USA

**Keywords:** biophysical modeling, transcription and translation, mammalian cell free expression, bacterial cell free expression

## Abstract

Cell-free expression (CFE) systems have been used extensively in system and synthetic biology as a promising platform for manufacturing proteins and chemicals. Currently, the most widely used CFE system is *in vitro* protein transcription and translation platform. As the rapidly increased applications and uses, it is crucial to have a standard biophysical model for quantitative studies of gene circuits, which will provide a fundamental understanding of basic working mechanisms of CFE systems. Current modeling approaches mainly focus on the characterization of *E. coli*-based CFE systems, a computational model that can be utilized to both bacterial- and mammalian-based CFE has not been investigated. Here, we developed a simple ODE (ordinary differential equation)-based biophysical model to simulate transcription and translation dynamics for both bacterial- and mammalian-based CFE systems. The key parameters were estimated and adjusted based on experimental results. We next tested four gene circuits to characterize kinetic dynamics of transcription and translation in *E. coli*- and HeLa-based CFE systems. The real-time transcription and translation were monitored using Broccoli aptamer, double stranded locked nucleic acid (dsLNA) probe and fluorescent protein. We demonstrated the difference of kinetic dynamics for transcription and translation in both systems, which will provide valuable information for quantitative genomic and proteomic studies. This simple biophysical model and the experimental data for both *E. coli*- and HeLa-based CFE will be useful for researchers that are interested in genetic engineering and CFE bio-manufacturing.

## Introduction

Cell-free expression (CFE) systems are a widely used tool in systems and synthetic biology for applications including synthesis of toxic proteins, rapid enzyme engineering, and bacteriophage production.[1, 2] Over the last decade, CEF systems have been adjusted and reshaped to respond to the significant increasing interests for constructing complex biochemical systems or synthetic cells *in vitro* through the execution of genetic information.[3–5] Recently, with microfluidic techniques, CFE systems have been proven to be a useful quantitative platform to recapitulate biological processes *in vitro* by expressing synthetic gene circuits.[6–9] Compared to living systems, CFE systems are more adjustable for observation and manipulation, hence allowing rapid tuning of reaction conditions. Various approaches have been developed to prepare transcription-capable and translation-capable extracts from different host model organisms, including *E. coli* extract-based platform and HeLa-based platform.[10–12] Numerous platforms are now commercially available or can be easily prepared in laboratories. [13–16] For example, an *E. coli* extract-based CFE system was developed by Noireaux et al. by activating endogenous transcription from nature sigma factors for rapid testing of synthetic circuits. myTXTL CFE (Daicel Arbor Biosciences), PURExpress (New England Biolabs), and Expressway (Thermo Fisher Scientific) are three commercialized *E. coli* extract-based protein expression systems. The kinetic dynamics of various gene circuits, including gates, oscillators, and repressors have been characterized using an *E. coli*-based CFE protein synthesis system.[15] Meanwhile, with the widespread applications of CFE platforms, mammalian CFE platforms have continuously attracted the interests of researchers due to its post- and co-translational modifications.[17, 18] Currently, CHO cells, HEK293 cells, and HeLa cells based CFE systems are also commercially available (SinoBiological, ThermoFisher). A HeLa-based protein synthesis platform is supplemented with translation regulator to enhance translation efficiency was developed by our group and other groups. [6, 19] This HeLa-based CFE platform has been utilized to express soluble and membrane protein in artificial cells.[20, 21]

In CFE systems, it is necessary to characterize the kinetic dynamics of protein synthesis yield for quantitative studies and optimization of gene circuits. Transcription and translation kinetics are two key parameters that govern the performance and effectiveness of CFE systems. Thus, a better understanding of mRNA and protein synthesis dynamics is crucial for the applications of cell-free bio-manufacturing. To detect mRNA and protein dynamics, several groups have developed various approaches. Jaffrey et al. developed broccoli and spinach RNA aptamers to detect RNA activities in *E. coli*-based CFE, bacteria, and mammalian cells. Upon binding small molecules in CFE solution or live cells, these RNA aptamers activate the fluorescence of fluorophores.[22–24] These RNA aptmaers were used to track mRNA dynamics in bacterial CFE systems.[25] Moreover, binary probe, molecular beacon, and double-strand locked nucleic acid (dsLNA) probe were developed to monitor mRNA dynamics in CFE systems. [6, 26, 27] Several groups have established biophysical models that can simulate kinetic dynamics of mRNA and protein synthesis.[27–29] Varner et al. developed an effective modeling method to understand the dynamics of transcription and translation using myTXTL toolkit.[28] Noireaux et al. modeled the dynamics of genetic circuits in *E. coli*-based CFE systems.[25, 29] However, previous studies of transcription and translation dynamics mainly focus on reactions in *E. coli*-based CFE systems. Hereby, a standard simple biophysical model is necessary to characterize transcription and translation kinetics in both bacterial- and mammalian-based CFE systems.

In this article, we developed a simple standard biophysical model to simulate the kinetic dynamics of transcription and translation in both *E. coli*-based and HeLa-based CFE systems. Meanwhile, we characterized the dynamics of mRNA and protein synthesis process by monitoring real-time synthesized mRNA and protein. Using a commercial *E.coli*-based CFE system (myTXTL toolkit, Daicel Arbor Bioscience), we simulated and experimental characterized a single promoter of Broccoli, a single promoter of deGFP, and a two-stage transcriptional cascade. Experimental studies were performed to evaluate the effectiveness of this simple biophysical model. In the HeLa-based CFE system, we utilized a dsLNA probe to monitor the real-time expression dynamics of synthesized mRNA. The synthesized protein dynamics were monitored using a GFP reporter. The kinetic dynamics of mRNA and protein expression were also modeled using this simple biophysical model. This model predicts that the synthesized protein is template DNA concentration dependent, and time- and resource-dependent. Our results demonstrated the ability of this simple biophysical model for the characterization of complex chemical reactions and biological parts. This simple biophysical model, together with dsLNA probe and broccoli aptamers, provide an efficient and versatile platform for characterizing transcription and translation dynamics in gene circuits in the context of artificial cells.

## 2. Materials and Methods

### 2.1 Preparation of *E. Coli* extract-CFE System

*E. coli*-based CFE system (myTXTL σ_70_ Master Mix) and all the plasmids were purchased from Daicel Arbor Biosciences (Ann Arbor, MI) in 1.5 mL centrifuge tubes. The sigma 70 master mix was thawed on ice for 10 minutes. The master mix is 75% of the final volume, the 25% left are for DNA and water. The DNA plasmids (pTXTL-P70a-broccoli, pTXTL-P70a-deGFP, pTXTL-P70a-S28, pTXTL-P28a-deGFP) were purchased in lyophilized form and purified using Miniprep Kit (Qiagen) using DH5 α cell lines. For pTXTL-P70a-broccoli, a 40 μM DFHBI-1T was added. The total volume of each reaction was prepared with a volume of 12 μL. The reaction solution was vortexed for 2-3 seconds to avoid bubbles and added to 384-well plate (Nunc^™^ MicroWell^™^ 384-Well Optical-Bottom Plates, 142761). A 10 μL of prepared reaction solution was added to each well. A plate seal (Nunc^™^ Sealing Tapes, 232701, ThermoFisher) was used to seal the well plate to keep the temperature inside the wells. All the samples were prepared in duplicate. All the fluorescence measurements were taken in clear-bottom polypropylene microplates using a fluorescence microplate reader (BioTek, Syngery 2). The fluorescence intensity was measured at the excitation and emission wavelength of 488/525 for GFP every three minutes at 29 °C.

### 2.2 GFP dsLNA probe design and preparation

The dsLNA probe consists of two pieces of nucleotide sequences, donor sequence and quencher sequence. The donor is a 21-base nucleotide sequence with alternating LNA/DNA monomers. The donor sequences were designed to detect GFP mRNA based on the minimum free energy secondary structure using RNAFold web server. The design principle has been reported previously.[6, 30–33] Briefly, the donor sequence is complementary to partial of the target mRNA sequence. After choosing the donor sequence, the binding affinity was optimized using NCBI Basic Local Alignment Search Tool (BLAST) database. A fluorophore (Texas Red) was labeled at the 5’ end of the donor sequence for fluorescence detection. The quencher is a 10-base LNA/DNA nucleotide sequence labeled with an Iowa Black RQ to quench the red fluorescence of donor. All the LNA probes and DNA sequences were synthesized by Integrated DNA Technologies (IDT).

To prepare dsLNA probe solution, donor and quencher were prepared in distilled water at a concentration of 100 nM. The donor and quencher were mixed at the ratio of 1:2 (volume ratio). The mixed solutions were incubated at 95°C in a PCR machine for 5 minutes and cooled down to room temperature over 4 hours. The prepared LNA donor and quencher sequence were then ready to use for mRNA detection in HeLa-based CFE reactions.

### 2.3 Preparation of HeLa-based cell free expression system

HeLa-based cell free expression system was prepared following the previously reported procedure [6, 7]. Briefly, the HeLa lysate was prepared from spinner cultured HeLa S3 cells using minimal essential medium eagle medium (eMEM). After 5-7 days of culture, cells were harvested, lysed, and aliquoted at the concentration of 2 x 10^5^ cells/mL. The rest components of HeLa-based cell free expression system include truncated GADD34 (stock concentration 2.3 mM), T7 RNA polymerase (stock concentration 5 mM), mix 1, and mix 2 solution. Mix 1 is a solution prepared with 27.6 mM Mg(OAc)2, 168 mM 4-(2-hydroxyethyl)-1-piperazineethanesulfonic acid (K-HEPES pH 7.5). Mix 2 is a solution which was prepared with the following reagents: 12.5 mM ATP, 8.36 mM GTP, 8.36 mM CTP, 8.36 mM UTP, 200 mM creatine phosphate, 7.8 mM K-HEPES (pH 7.5), 0.6 mg/mL creatine kinase. All these solutions were prepared based on previously reported protocol.[6] The HeLa-based cell free expression reactions were prepared as follows: first, HeLa lysate, GADD34, and mix solution were mixed, vortexed, and incubated in a PCR machine at 32°C for 15 minutes. The rest components were mixed together. For each component, the following volume was used: lysate (4.5 μL), GADD34 (1.35 μL), T7 (0.9 μL), mix1 (1.125 μL) and mix2 (1.125 μL). The DNA plasmids and water have a total volume of 3 μL. The samples were prepared in duplicate with a final volume of 12 μL per sample. A 10 μL of prepared reaction solution was added to each well. The pT7-CFE-GFP plasmid was purchased from Thermo Fisher Scientific and prepared at different concentrations.

### 2.4 Calibration of GFP standard curve

Recombinant GFP (MW, 29 kDa) was acquired from Cell BioLabs and prepared in 1x PBS with a concentration of 1 mg/mL. To generate a GFP standard calibration curve, a serial dilution of GFP was performed in a solution containing 12.5 mM Tris-HCl, 150 mM NaCl, and 50% glycerol. The serial dilution solutions were prepared using a 384 well plate. The fluorescence intensity of GFP with differentiation concentrations was measured in a microplate reader using excitation and emission wavelengths of 488 and 525 nm, respectively. The gain was set to 100 for all the experiments. Both serial dilutions and fluorescence calibration were performed in triplicate, and the average fluorescence reading for each concentration was obtained. Linear regression was used to estimate absolute GFP concentrations in CFE experiments.

### 2.4 DNA quantification using nanodrop

All the DNA plasmids were quantified using a nanodrop. First, a 2 μL of blank solution (distilled water) was added and calibrated as ‘Blank’. Then load DNA solution (2 μL) and measure the nucleic acid concentration as ng/μL. Each plasmid was measured twice and the average was obtained. The concentration was then converted to nM based on the average mass of one DNA base pair is 650 g/mol.

### 2.5 Biophysical model for translation and translation dynamics

A simple biophysical model was developed to study the dynamics of transcription (mRNA) and translation (protein) in bacterial- and mammalian-based CFE systems. Ordinary differentiation equations were used to model the mRNA levels and protein levels as they change with time (the rate change of mRNA and protein). In these differentiation equations, positive terms describe how chemical species are produced; negative terms describe how it degrades or is removed. In this model, mRNA levels for a gene expressed from a constitutive promoter have constant production, and mRNA degradation is proportional to the amount of mRNA present. To model translation, the mRNA is first translated into immature protein which will fold and form mature functional protein, **Figure S1**. Thus, the kinetics of transcription and translation can be described using the following differentiation equations (1–3).

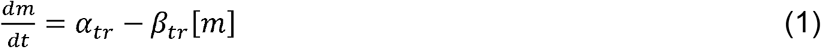

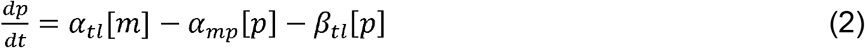

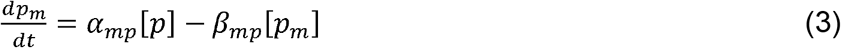

In this model, the rate change of mRNA, immature protein, and mature protein can be described by *dm*/*dt*, *dp*/*dt*, and *dp_m_*/*dt*, respectively. Here, [*m*] is the concentration of transcribed mRNA, [*p*] is the concentration of translated immature protein, [*p_m_*] is the concentration of translated mature protein. *α_tr_*, *α_tl_* and *α_mp_* are the first order transcription rate, translation rate, and protein maturation rate, respectively. The degradation rate of mRNA, immature protein, and mature protein are described using *β_tr_*, *β_tl_*, and *β_mp_*, respectively. In our experimental systems, there is no degradation observed, thus the degradation rates are set to zero (*β_tr_*= *β_tl_* = *β_mp_* =0). As indicated in equation (1), the mRNA level ([*m*]) depends on the transcription rate. The synthesized protein level depends on the number of mRNAs, and translation rate. The transcription rate and translation rate can be determined based on experimental data. The transcribed mRNA and translated mature protein levels were first quantified by measuring average fluorescence intensity. The simulation curves were then fitted by adjusting the transcription rate *β_tr_* and translation rate *β_tl_*.

### 2.6 Statistical analysis

Data are presented as mean ± s.e.m. All the measurements were conducted in duplicate, and repeated at least three independent times. Student’s *t*-tests were performed to analyze statistical significance between experimental groups. Statistically significant *p* values were assigned as follows: *, *p*<0.05; **, *p*<0.01 or ***, *p*<0.001.

## 3 Results

### 3.1 Modeling and characterization of kinetic dynamics of RNA regulation in *E. coli*-based CFE system

We first utilized this simple biophysical model to study the kinetic dynamics of transcription in *E. coli*-based CFE systems using sigma factor 70 (σ_70_) induced broccoli expression, **Figure 1A**. It has been reported that Broccoli and Spinach are RNA aptamers that bind to GFP fluorophore (DFHBI) and switch on the fluorescence.[22, 34] Broccoli is a 49-nt-long aptamer that was developed based on SELEX protocol that exhibits bright green fluorescence upon binding DFHBI or the improved version of this fluorophore, (Z)-4-(3,5-difluoro-4-hydroxybenzylidene)-2-methyl-1-(2,2,2-trifluoroethyl)- 1H-imidazol-5(4H)-one) (DFHBI-1T) (DFHBI-1T).[35] It has been reported that RNA aptamers can be added to CFE reactions (like Broccoli or Spinach aptamer) to monitor the dynamics of mRNA synthesis.[25] RNA aptamers bind to specific dyes, which only fluoresce when they are bound to the mRNA. To express fluorescent RNA aptamers, we prepared the reaction solutions based on manufacturing’s instructions. Then broccoli aptamer DNA template (pTXTL-P70a-Broccoli) at different concentrations (0, 0.5, 1, 2, 5, 10 nM) were added. The corresponding fluorescent dye, DFHBI-1T were added. To make sure the concentration of the dye is always in excess to synthesized RNA, 40 μM of DFHB1-1T was added for Broccoli RNA aptamer. The transcription dynamics is quantified by measuring fluorescence intensity with the excitation/emission wavelength at 488/525 nm every three minutes for two hours. The synthesized mRNA concentrations were calculated according to the calibration curve, **Figure S2.** Furthermore, we examined synthesized mRNA concentrations at different DNA concentrations. After two hours of incubation, the synthesized mRNA obtained were 0.08 μM, 0.32 μM, 0.97 μM, 3.54 μM, and 10.1 μM at DNA concentrations of 0.5 nM, 1 nM, 2 nM, 5 nM, and 10 nM, respectively, **Figure 1C**. The mRNA expression dynamics were modeled using our model. The synthesized mRNA concentrations versus time were plotted, **Figure 2B**. The kinetic constants were estimated based on literation and our experimental results, **Table S2**. The modeling results showed a similar profile of mRNA synthesis dynamics compared to our experimental results. We next calculated mRNA production rate: [mRNA] production rate [μM/min] = (fluorescence intensity at current time point – fluorescence intensity at previous time point) / time interval. For single promoter regulations, the production rate of ‘broccoli’ mRNA was calculated and compared, **Figure 1D**. The results indicate that synthesized mRNA was produced continuously over the first two hours of the period. The ‘broccoli’ production rate increased in the first 30 minutes and decreased sharply afterward due to mRNA degradation. After two hours, the synthesized mRNA concentration reached to peak, with the lowest mRNA production rate (about 0 μM/min).

**Figure 1.**
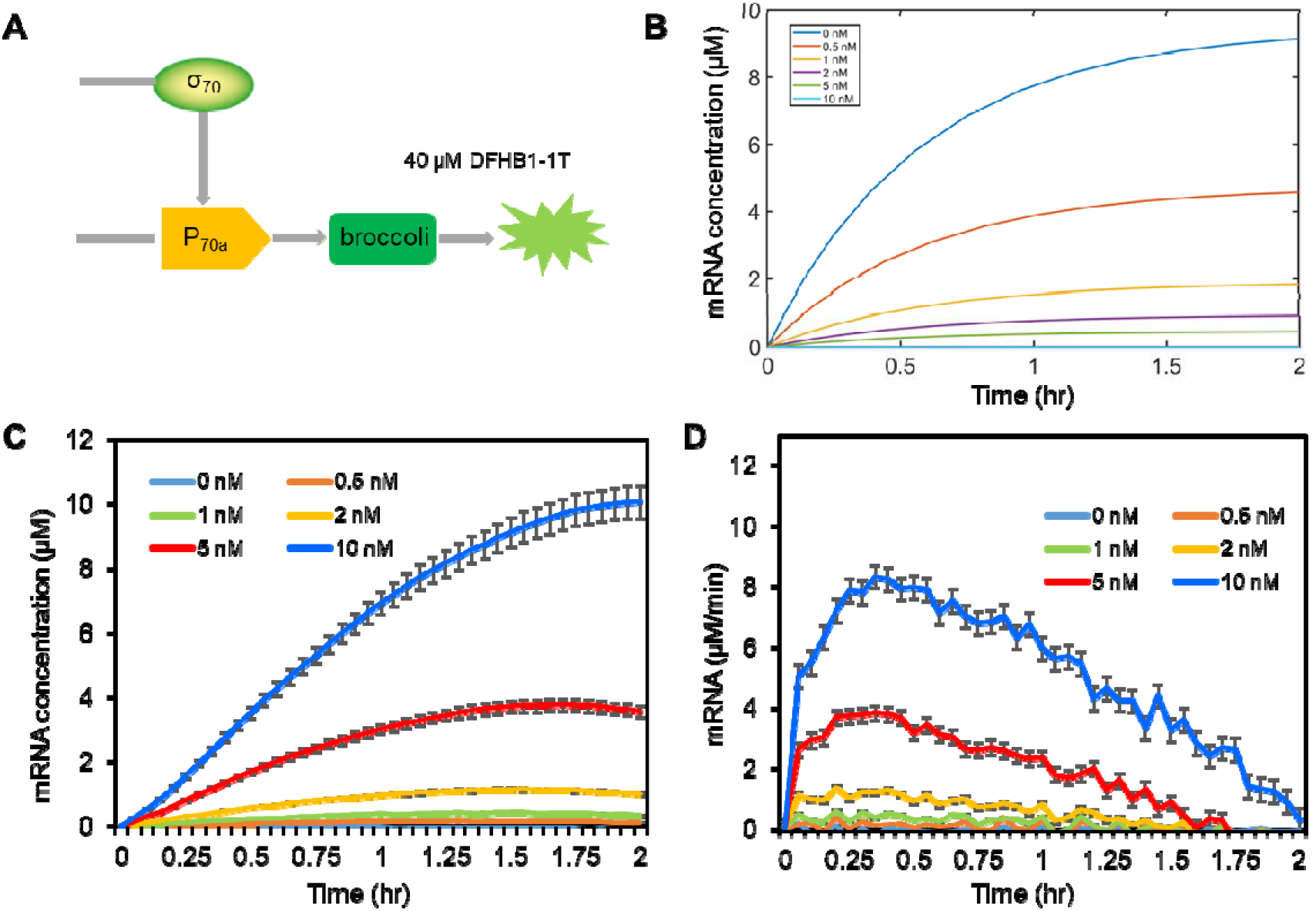
Modeling and characterization of transcription dynamics of a single promotor of myTXTL-P70a-Broccoli in *E.coli*-based CFE. **(A)** Schematic illustration of a single promotor of P70-Broccoli. **(B)** Modeling results of mRNA expression dynamics at different DNA concentrations. The transcription rate was estimated based on reference and fitted using our experimental results. **(C)** Kinetic dynamics of mRNA synthesis in CFE at different plasmid DNA concentrations of 0.5 nM, 1 nM, 2 nM, 5 nM, and 10 nM. A negative control was designed when there was no DNA plasmid added. **(D)** mRNA production rate at different DNA plasmid concentrations. Experiments were repeated at least three times independently (n=5). Data are expressed as mean ± s.e.m.

**Figure 2.**
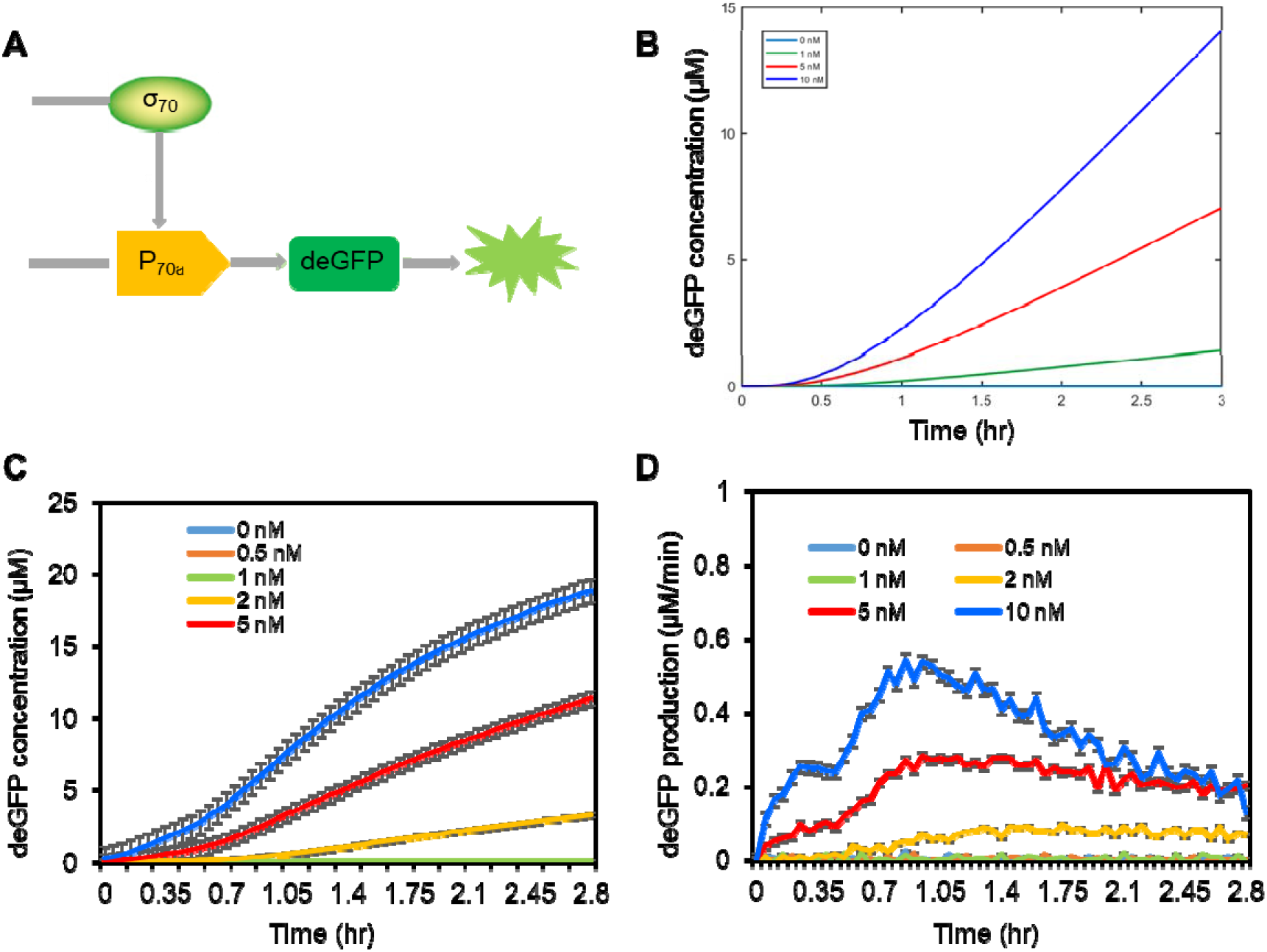
Modeling and characterization of translation dynamics of a single promotor of myTXTL-P70-deGFP in *E.coli*-based CFE. **(A)** Schematic illustration of single promoter of myTXTL-P70-deGFP. **(B)** Modeling results of protein expression dynamics at different DNA concentrations. The transcription and translation rate was estimated based on reference and fitted using our experimental results. **(C)** Kinetic dynamics of protein synthesis in CFE at plasmid DNA (mTXTL-P70-deGFP) concentrations of 0.5 nM, 1 nM, 2 nM, 5 nM, and 10 nM. **(D)** Protein production rate in CFE at different DNA plasmid concentrations. Experiments were repeated at least three times independently (n=7). Data are expressed as mean ± s.e.m.

### 3.2 Modeling and characterization of kinetic dynamics of protein expression in *E. coli*-based CFE system

We next investigated the kinetic dynamics of protein synthesis in an *E. coli*-based CFE system using two circuits. One is single promotor p70-deGFP, the other one is a two-stage transcriptional activation cascade using sigma 28 (σ_28_). Both circuits start with a specific sigma promotor, sigma 70, which is present in the cytoplasmic extract. The translation were first modeled and characterized with a single promotor p70-deGFP, **Figure 2A**. To model the kinetic dynamics of translation, the simple biophysical model was utilized, including equation (1-3). The synthesized protein concentrations were modeled with different plasmid DNA concentrations, ranging from 1 nM to 10 nM, **Figure 2B**. Based on this simple biophysical model, the synthesized protein concentrations with the DNA concentrations of 1 nM, 5 nM, and 10 nM were 1.5 μM, 6.7 μM, and 14.8 μM. We further characterized protein synthesis dynamics in the *E.coli*-based CFE system. **Figure 2C** shows protein expression kinetics with different p70a-deGFP concentrations, ranging from 0.5 nM to 10 nM. A negative control was used when there were no DNA plasmids added. The protein production increases within the first three hours. Protein synthesis as a function of DNA template is only linear with the DNA concentration from 1 nM to 5 nM, **Figure S5**. Without DNA plasmids, the protein synthesis is negligible. The production rate was calculated as [deGFP] production rate [μM/min] = (fluorescence intensity at current time point – fluorescence intensity at previous time point)/time interval. The fluorescence intensity was then converted to μM based on the calibration curve, **Figure S3**. The production rate versus time was plotted, as shown in **Figure 2D**. The protein production rate increased in the first hour and slowed down gradually, indicating that the protein synthesis depends on the DNA concentrations and resources in the master mix. It has been reported that gene expression in the CFE system is independent of the resources only for a short period of time (1~2 hours for conventional systems).[13] The kinetics of gene expression can be altered by the decrease of the energy charge, degradation of amino acids, and pH change during transcription and translation reactions.

This simple biophysical model was further utilized to simulate a two-stage transcriptional activation cascade with *E. coli* sigma 28 (σ_28_) transcription factor, as illustrated in **Figure 3A**. Transcriptional activation cascades are simple gene circuits that require expression of a transcription factor, including sigma 19, 24, 28, 54, and 70, to activate the expression of fluorescent protein (deGFP, mCherry, or RFP). [36] Here, in our circuit, there are two steps of transcription and translation. We first need to activate the expression of sigma factor 28, which is required to activate the next transcription and translation process to produce deGFP, **Figure 3A**. To model this two-stage transcriptional activation cascade, we developed six differentiation equations based on our simple model, (**Supplementary Information**). **Figure 3B** shows the modeling results of synthesized protein expression in transcriptional activation cascade based on the different DNA concentrations ranging from 1 nM to 10 nM. Based on these two-stage transcription translation processes, the synthesized protein in this transcription activation cascade are 1.1 nM, 2.02 nM, 5.5 nM, and 11 nM with the DNA concentrations of 1 nM, 2 nM, 5 nM, and 10 nM, respectively. The kinetic dynamics of protein synthesis in this two-stage cascade showed a translation delay due to the generation of sigma 28, which is required for the activation of deGFP. Next, we characterized this cascade using *E. coli*-based CFE systems. In this cascade reaction, there are two DNA plasmids, pTXTL-P70a-S28, and pTXTL-P28a-deGFP. Here the concentration of pTXTL-P70a-S28 was set to 0.05 nM for comparison. By adjusting the DNA (pTXTL-P28a-deGFP) concentrations (0.5 nM, 1 nM, 2 nM, 5 nM, and 10 nM), the dynamics of synthesized deGFP were monitored over the period of ~3 hours with an interval of 3 minutes. A negative control was designed when there was no DNA presented. The fluorescence intensity of synthesized deGFP were measured using excitation and emission wavelengths of 488 and 525 nm, respectively. The measured fluorescence intensity was then converted to deGFP concentrations based on the calibration curve, **Figure S3**. With the DNA concentrations of 0.5 nM, 1 nM, 2 nM, 5 nM, and 10 nM, the synthesized deGFP are calculated as 0.25 nM, 0.34 nM, 1.2 nM, 7.2 nM, and 11.5 nM, respectively. The production rate was calculated as [deGFP] production rate [μM/min] = (intensity at current time point – intensity at previous time point)/time interval. The production rate dynamics with different DNA concentrations were plotted, **Figure 3D**. The production rate was increased gradually in the first hour and was slowed down for the rest of the reaction period. In this two-stage transcription activation cascade, there are two limiting factors for the generation of deGFP, one is the pTXTL-P70a-S28 DNA concentration, and the other one is pTXTL-P28-deGFP concentration. The highest production rate can reach 0.3 μM/min with a DNA concentration of 10 nM. The kinetic dynamics of deGFP expression of this two-stage transcription translation cascade are similar to single promoter deGFP protein synthesis dynamics. A 10-30 minutes delay was observed for sigma 28 transcriptional activation cascaded compared to the expression of deGFP from a signal promoter P_70_, a typical time for such two-stage cascades. [37]

**Figure 3.**
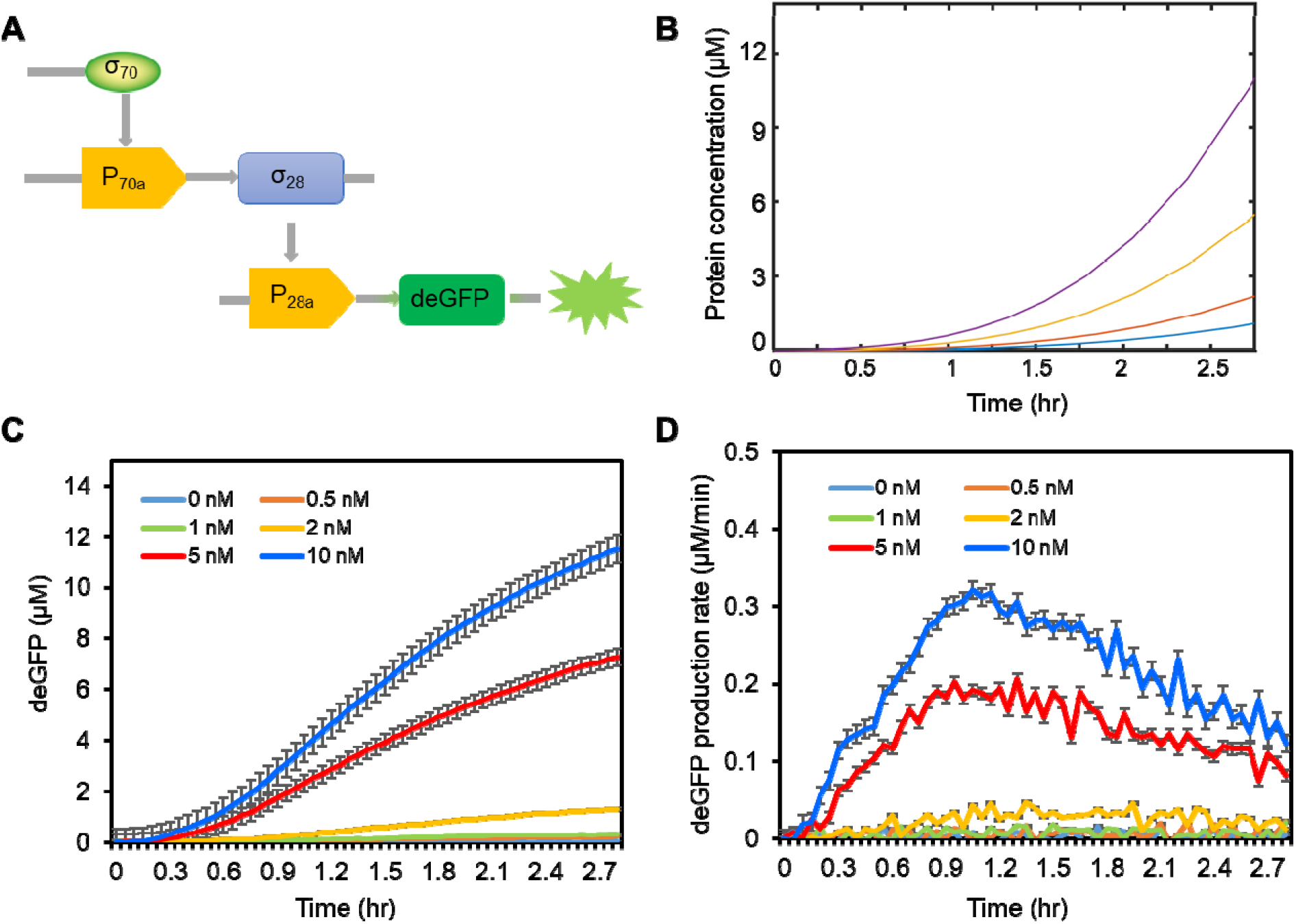
Modeling and characterization of translation dynamics of a two-stage cascade in *E.coli*-based CFE. **(A)** Schematic illustration of a two-stage transcriptional activation cascade. **(B)** Modeling results of protein expression dynamics at different DNA concentrations in a two-stage cascade. **(C)** Kinetic dynamics of deGFP synthesis in CFE at different plasmid DNA (pTXTL-P28a-deGFP) concentrations. A negative control was designed when there was no DNA plasmid added. **(D)** deGFP production rate at different DNA (pTXTL-P28a-deGFP) concentrations. The concentration of pTXTL-P70a-S28 was set to 0.05 nM. Experiments were repeated at least three times independently (n=5). Data are expressed as mean ± s.e.m.

### 3.3 Modeling and characterization of kinetic dynamics of RNA and protein in mammalian CFE system

Although bacterial CFE systems offer broad versatility, scalability and portability to study transcription and translation dynamics in different gene circuits, eukaryotic CFE systems are more advantageous due to their ability to carry out post-translational modifications.[20, 27, 38] Thus, it is important to understand the kinetic dynamics of transcription and translation in eukaryotic CFE systems. Recently, there is increasing evidence that HeLa cell-derived *in vitro* coupled transcription/translation system with supplemented transcription and translation factors plays an important role in bottom-up synthetic biology, including building synthetic cells. [6, 9, 39, 40] Thus, we utilized a HeLa-based CFE system to characterize transcription and translation dynamics. **Figure 4A** shows the illustration of a HeLa-based CFE system including different components. This HeLa-based CFE system allows an efficient means of producing any proteins of interest. The dynamics of synthesized protein were monitored using GFP reporter gene. To characterize the kinetic dynamics of transcription in this mammalian CFE system, we utilize a dsLNA probe to monitor the transcription dynamics, **Figure 4B**. Here, the dsLNA probe was designed to detect GFP mRNA in this HeLa-based CFE system. To prepare the reaction solutions, HeLa-based CFE solution was first prepared including HeLa lysate, truncated GADD34, T7, mix solution 1, and mix solution 2. The dsLNA and DNA plasmid were then added and incubated at 32 °C for 4 hours. The final concentration of dsLNA probe was set to 100 nM, which is sufficient to detect GFP mRNA as high as the concentration of 100 μM. The fluorescence intensity of synthesized mRNA and protein were measured with the wavelength of excitation and emission at 590/617 (red channel) and 488/525 (green channel), respectively, **Figure 4C**.

**Figure 4.**
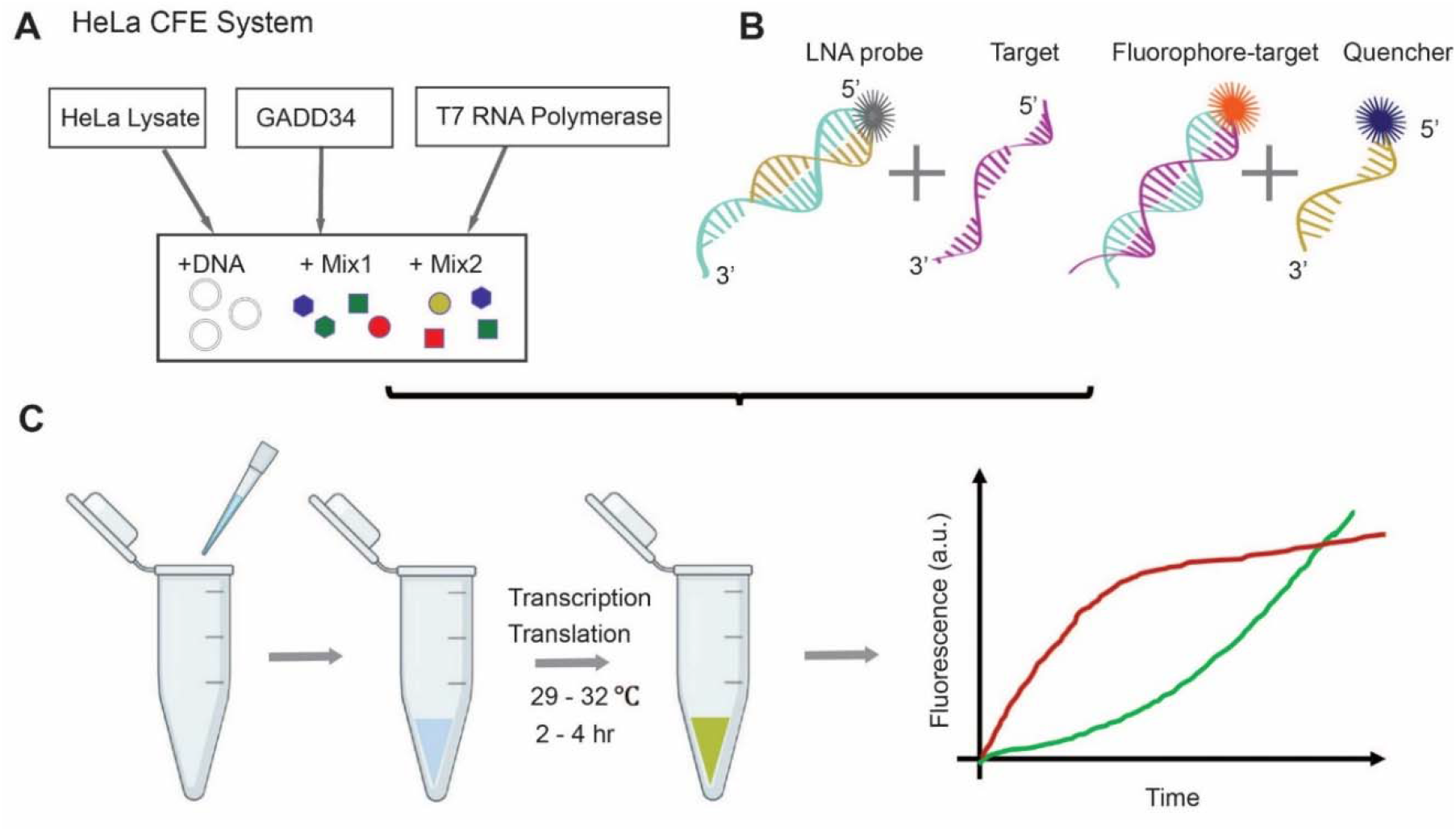
Illustration of real-time detection of transcription and translation dynamics using HeLa-based CFE system and dsLNA probe. **(A)** Components of HeLa-based CFE system. **(B)** Working principle of dsLNA probe for mRNA detection. **(C)** Illustration of the experimental setup for monitoring transcription and translation dynamics in HeLa-based CFE.

To characterize kinetic dynamics of transcription and translation in HeLa-based CFE, the simple biophysical model with transcription, translation, and maturation was utilized to model the process, equation (1-3). **Figure 5A** and **Figure 5B** shows the modeled GFP mRNA and protein concentrations with different DNA concentrations ranging from 1 nM to 5 nM, respectively. The mRNA and protein synthesis rates were estimated based on experimental results, **Supplementary Table S2**. The profile of synthesized mRNA is referred to hyperbolic and demonstrated saturation after one hour due to the limited resources and energy in the CFE reaction solution. Meanwhile, the protein synthesis process showed an S-shaped curve (sigmoid curve), indicating the slow protein synthesis and maturation process.

**Figure 5.**
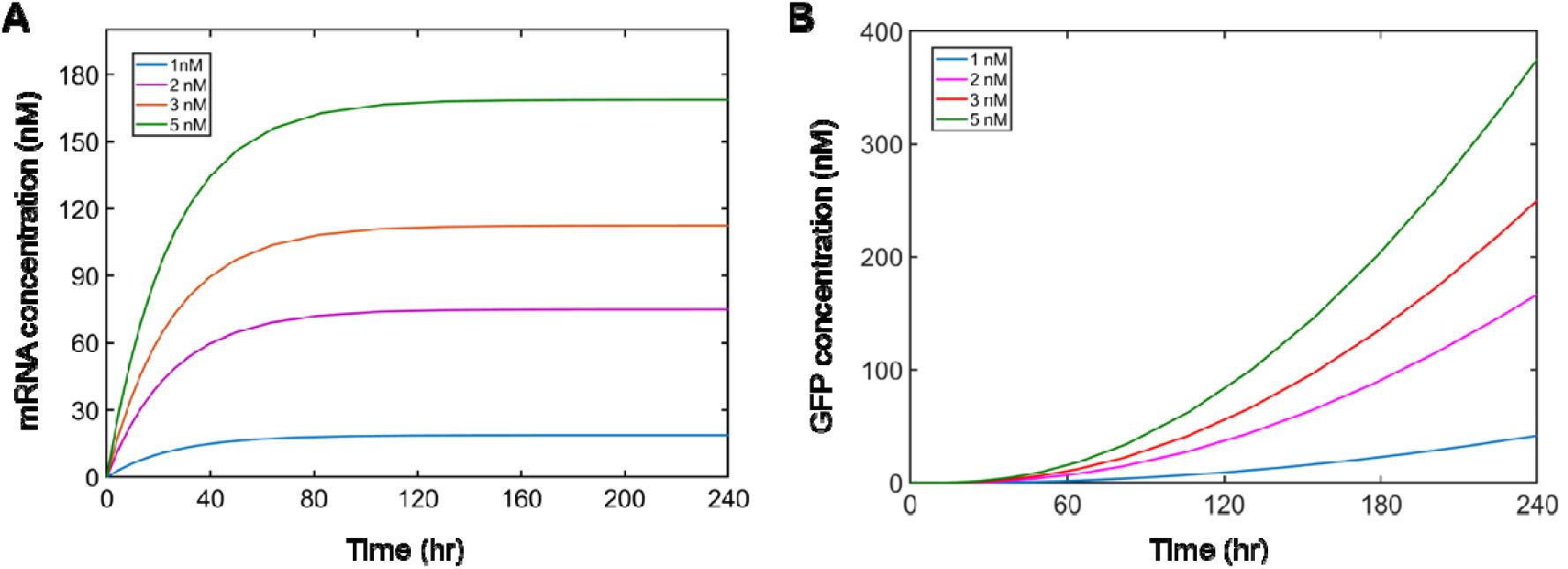
Modeling results of transcription and translation dynamics in HeLa-based CFE. **(A)** Simulation results of mRNA synthesis in CFE at different DNA concentrations of 1 nM, 2 nM, 3 nM, and 5 nM. **(B)** Simulation results of protein synthesis in CFE at different DNA concentrations of 1 nM, 2 nM, 3 nM, and 5 nM.

To further characterize the kinetic dynamics of transcription and translation in HeLa-based CFE, the mRNA and protein expression levels were monitored on a fluorescence plate reader over 4 hours with sampling every 3 minutes. The transcription and translation dynamics of HeLa-based CFE were quantified by measuring fluorescence intensity following the respective excitation/emission wavelengths. The synthesized mRNA and protein concentrations were calculated according to the dsLNA probe and GFP calibration curves, respectively, **Figure S3** and **S4**. The mRNA and protein expression dynamics over 4 hours of incubation period were plotted, **Figure 6A**. The synthesized mRNA concentrations obtained were 20.2 nM, 51.7 nM, 109.2 nM, and 159.5 nM at DNA concentrations of 1 nM, 2 nM, 3 nM, and 5 nM, respectively. The synthesized protein concentrations obtained were 30.2 nM, 90.2 nM, 161.4 nM, and 218.7 nM at DNA concentrations of 1 nM, 2 nM, 3 nM, and 5 nM, respectively, **Figure 6A** and **6C**. These results indicate that GFP mRNA was produced and could be detected immediately by the dsLNA probe, while GFP protein was not detected until almost 60 minutes due to GFP maturation. Unlike *E.coli* – based CFE system, the monotonic increase of mRNA and protein in HeLa-based CFE systems were non-linear with respect to both time and DNA concentration, **Figure S6**. The origins of these differences are not clear; however, the results show the high need for extensive characterization of mRNA and protein expression dynamics in mammalian CFE systems, which may have different mechanisms compared to bacterial CFE systems.

**Figure 6.**
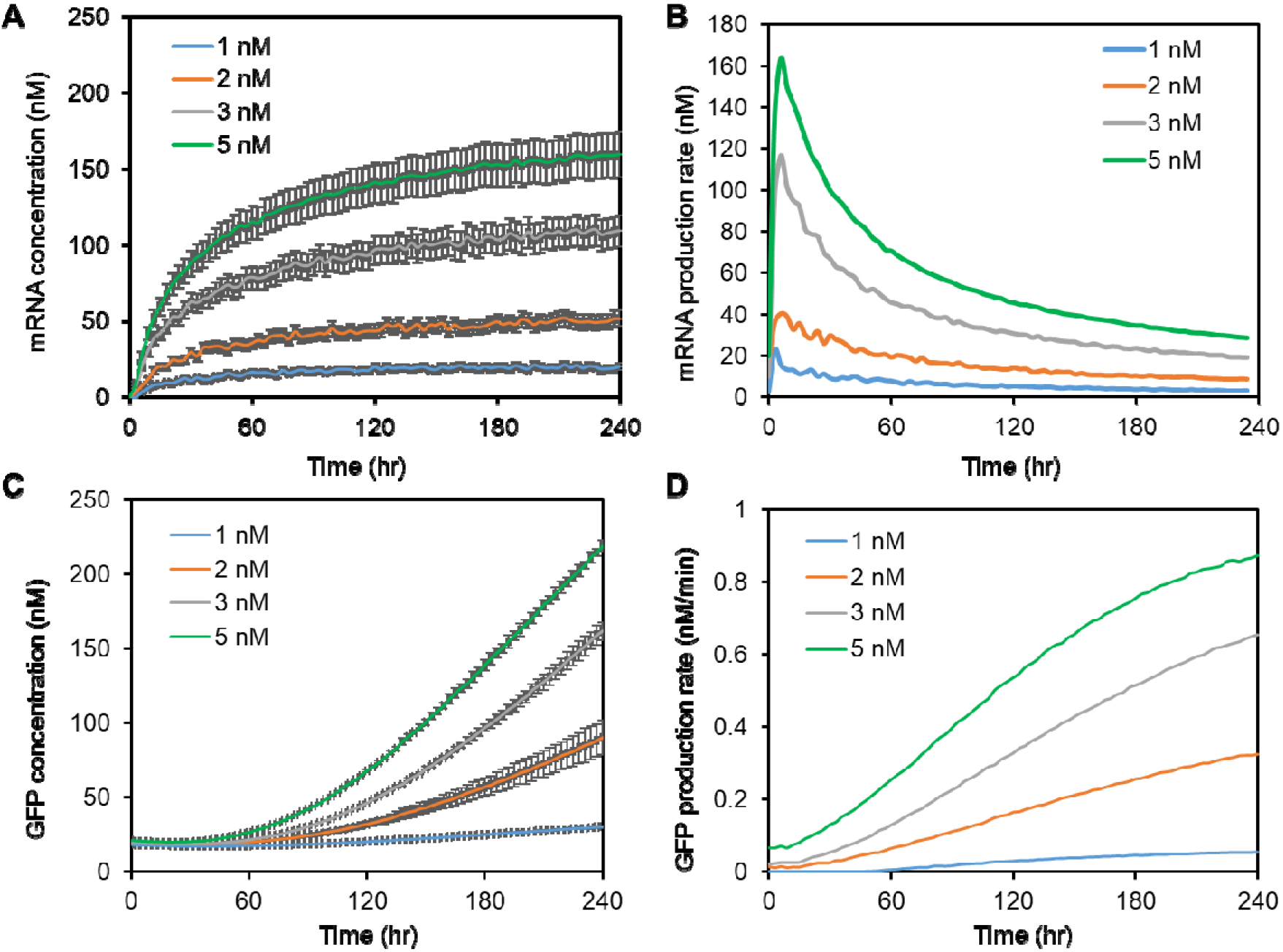
Characterization of mRNA and protein synthesis dynamics in HeLa-based CFE. **(A)** mRNA synthesis dynamics at different plasmid DNA concentrations. **(B)** mRNA production rate in CFE at different DNA concentrations. **(C)** Protein synthesis dynamics at different plasmid DNA concentrations. **(D)** Protein production rate in CFE at different DNA concentrations. The DNA (pT7-CFE-GFP) concentrations were set to 1 nM, 2 nM, 3 nM, and 5 nM, respectively. The LNA probe was 100 nM for all the experiments. Experiments were repeated three times independently. Data are expressed as mean ± s.e.m.

The mRNA and protein production rate were calculated as [nM/min] = (mRNA or protein concentration at current time point – mRNA or protein concentration at previous time point)/ time interval. The mRNA and protein production rate dynamics were then plotted, **Figure 6B** and **6D**. The mRNA production rate reached to its peak after about 10 minutes of incubation and decreased sharply after 30 minutes of reaction. The highest production rate was measured at 163.8 nM/min with a DNA concentration of 5 nM. It is noted that the mRNA expression dynamics observed here may be specific to HeLa-based CFE systems utilizing T7 RNA polymerase, which increased transcription rate substantially relative to endogenous transcription machinery. Compared to mRNA, the synthesized protein was increased slowly and can be continuously expressed for ~ 10 hours (**Supplemental Figure S7**). The highest production rate of protein is 0.91 nM/min with a DNA concentration of 5 nM after 4 hours of incubation. These results indicated that in mammalian CFE systems, transcription and translation are separated in time and follow different dynamics.

## Discussion

Cell free gene expression systems originally were developed as a tool for quick protein synthesis. Over the last decade, emerging evidence showed that CFE systems are important for high-throughput expression screening, high yield protein production, synthetic and systems biology applications.[2, 4, 38, 39] Recently, CFE system was used widely as an experimental platform for bottom-up synthetic biology to build artificial cells. [41, 42] Transcription and translation are two important processes that govern metabolism and signal transduction. However, CFE systems from different origins (bacterial, mammalian) may have different dynamics in terms of reaction speed, expected yield, or kinetic parameters. In this article, we established an approach to characterize the mRNA and protein synthesis processes in both *E.coli*–based CFE and HeLa-based CFE systems. A simple biophysical model was developed to simulate the kinetic dynamics of transcription and translation processes in CFE systems. For *E.coli*-based CFE, three gene circuits, including single promoter of P70-Broccoli, single promoter of P70-deGFP, and transcriptional activation cascade were tested and compared. For HeLa-based CFE, the transcription and translation were characterized using dsLNA probe and pT7-CFE-GFP. It is noted that although this simple biophysical model can be adjusted for all the transcription and translation dynamics in CFE systems, the reaction speed, including transcription and translation rate, degradation rate may be quite different for different CFE systems.

The *E. coli*-based CFE systems are formed with sigma factor 70 (σ_70_). There are seven native transcription factors to E. coli: σ19, σ24, σ28, σ32, σ38, σ54, and σ70. Each sigma factor is expressed in *E. coli* in response to different conditions.[43] The sigma promoter P70a, originates from the lambda phage repressor Cro promoter, is the housekeeping sigma factor and is responsible for expressing most genes in *E. coli.*This *E. coli* sigma 70 promoter is the strongest promoter so far reported.[44] For single promotor P70-Broccolli and P70-deGFP, the mRNA expression was detected immediately using Broccoli aptamer with minimum delay, the deGFP protein expression was detected as early as several minutes after the reactions started. These results indicate σ70 is a strong promoter to drive the transcriptional regulations presented in *E. coli.* For a two-stage transcriptional cascade, there is a 30 minutes delay for protein expression. It was also observed that the delay is larger for σ_28_ transcriptional activation units at low plasmid concentrations (i.e., 0.5 nM, 1 nM). This could be potentially caused by the high efficiency of proteolysis with the SsrA tag. [13, 45] Another limiting factor for CFE systems is the availability of necessary resources, especially for slow processes requiring a significant amount of energy for transcription and translation. The HeLa-based CFE system is formed using T7 RNA polymerase due to its widespread adoption of T7 promoter in many CFE applications. T7 bacteriophage promoter allows *in vitro* transcription as strong as *in vivo* conditions. In our HeLa-based CFE system, the synthesis of mRNA starts right after the reaction starts without delay, while the process of protein synthesis has about 1-hour delay due to GFP maturation. These results demonstrated that bacterial CFE and mammalian CFE systems follow different transcription and translation dynamics.

Although there are a variety of commercial cell-free expression systems (i.e., PURExpress, myTXTL, and 1-step human high yield IVT), the dynamics of transcription and translation were rarely characterized and simulated at the same time. Here, for the first time, we characterized the kinetic dynamics of transcription and translation in *E. coli*-based and HeLa-based CFE systems, using Broccoli aptamer, dsLNA probe and fluorescent protein. For the HeLa-based CFE system, we simultaneously monitored the transcription and translation dynamics. We demonstrated the difference of kinetic dynamics for transcription and translation in both systems, which will provide valuable information for quantitative genomic and proteomic studies. With proper characterization and quantitative biophysical modeling of *in vitro* expression kinetics will eventually turn both bacterial and mammalian CFE systems into a versatile tool for synthetic biology and systems biology.

## Acknowledgement

The authors would like to thank Cold Spring Harbor Laboratory (CSHL) Synthetic Biology (Synbio) Summer Course. This work is supported by the NASA CT SGC #P-1558 to S. Wang.

## Notes

### Competing Interest Statement

The authors have declared no competing interest.

## References

[1] M. B. Kopniczky, C. Canavan, D. W. McClymont, M. A. Crone, L. Suckling, B. Goetzmann, et al., “Cell-free protein synthesis as a prototyping platform for mammalian synthetic biology,” ACS synthetic biology, vol. 9, pp. 144–156, 2020.

[2] A. D. Silverman, A. S. Karim, and M. C. Jewett, “Cell-free gene expression: an expanded repertoire of applications,” Nature Reviews Genetics, vol. 21, pp. 151–170, 2020.

[3] V. Noireaux and A. P. Liu, “The new age of cell-free biology,” Annual review of biomedical engineering, vol. 22, pp. 51–77, 2020.

[4] M. T. Smith, K. M. Wilding, J. M. Hunt, A. M. Bennett, and B. C. Bundy, “The emerging age of cell-free synthetic biology,” FEBS letters, vol. 588, pp. 2755–2761, 2014.

[5] D. Garenne and V. Noireaux, “Cell-free transcription–translation: engineering biology from the nanometer to the millimeter scale,” Current opinion in biotechnology, vol. 58, pp. 19–27, 2019.

[6] S. Wang, S. Majumder, N. J. Emery, and A. P. Liu, “Simultaneous monitoring of transcription and translation in mammalian cell-free expression in bulk and in cell-sized droplets,” Synthetic Biology, vol. 3, p. ysy005, 2018.

[7] K. K. Ho, V. L. Murray, and A. P. Liu, “Engineering artificial cells by combining HeLa-based cell-free expression and ultrathin double emulsion template,” Methods in cell biology, vol. 128, pp. 303–318, 2015.

[8] K. K. Ho, L. M. Lee, and A. P. Liu, “Mechanically activated artificial cell by using microfluidics,” Scientific reports, vol. 6, pp. 1–10, 2016.

[9] S. Majumder and A. P. Liu, “Bottom-up synthetic biology: modular design for making artificial platelets,” Physical biology, vol. 15, p. 013001, 2017.

[10] J. G. Perez, J. C. Stark, and M. C. Jewett, “Cell-free synthetic biology: engineering beyond the cell,” Cold Spring Harbor perspectives in biology, vol. 8, p. a023853, 2016.

[11] B. C. Bundy, J. P. Hunt, M. C. Jewett, J. R. Swartz, D. W. Wood, D. D. Frey, et al., “Cell-free biomanufacturing,” Current opinion in chemical engineering, vol. 22, pp. 177–183, 2018.

[12] T. Kigawa, T. Yabuki, N. Matsuda, T. Matsuda, R. Nakajima, A. Tanaka, et al., “Preparation of Escherichia coli cell extract for highly productive cell-free protein expression,” Journal of structural and functional genomics, vol. 5, pp. 63–68, 2004.

[13] J. Shin and V. Noireaux, “An E. coli cell-free expression toolbox: application to synthetic gene circuits and artificial cells,” ACS synthetic biology, vol. 1, pp. 29–41, 2012.

[14] D. Garenne, S. Thompson, A. Brisson, A. Khakimzhan, and V. Noireaux, “The all-E. coliTXTL toolbox 3.0: new capabilities of a cell-free synthetic biology platform,” Synthetic Biology, vol. 6, p. ysab017, 2021.

[15] J. Garamella, R. Marshall, M. Rustad, and V. Noireaux, “The all E. coli TX-TL toolbox 2.0: a platform for cell-free synthetic biology,” ACS synthetic biology, vol. 5, pp. 344–355, 2016.

[16] R. W. Martin, N. I. Majewska, C. X. Chen, T. E. Albanetti, R. B. C. Jimenez, A. E. Schmelzer, et al., “Development of a CHO-based cell-free platform for synthesis of active monoclonal antibodies,” ACS synthetic biology, vol. 6, pp. 1370–1379, 2017.

[17] L. Thoring, D. A. Wüstenhagen, M. Borowiak, M. Stech, A. Sonnabend, and S. Kubick, “Cell-free systems based on CHO cell lysates: Optimization strategies, synthesis of “difficult-to-express” proteins and future perspectives,” PLoS One, vol. 11, p. e0163670, 2016.

[18] E. D. Carlson, R. Gan, C. E. Hodgman, and M. C. Jewett, “Cell-free protein synthesis: applications come of age,” Biotechnology advances, vol. 30, pp. 1185–1194, 2012.

[19] S. Mikami, M. Masutani, N. Sonenberg, S. Yokoyama, and H. Imataka, “An efficient mammalian cell-free translation system supplemented with translation factors,” Protein expression and purification, vol. 46, pp. 348–357, 2006.

[20] K. K. Ho, J. W. Lee, G. Durand, S. Majumder, and A. P. Liu, “Protein aggregation with poly (vinyl) alcohol surfactant reduces double emulsion-encapsulated mammalian cell-free expression,” PLoS One, vol. 12, p. e0174689, 2017.

[21] S. Majumder, N. Wubshet, and A. P. Liu, “Encapsulation of complex solutions using droplet microfluidics towards the synthesis of artificial cells,” Journal of Micromechanics and Microengineering, vol. 29, p. 083001, 2019.

[22] G. S. Filonov, J. D. Moon, N. Svensen, and S. R. Jaffrey, “Broccoli: rapid selection of an RNA mimic of green fluorescent protein by fluorescence-based selection and directed evolution,” Journal of the American Chemical Society, vol. 136, pp. 16299–16308, 2014.

[23] T. A. Rogers, G. E. Andrews, L. Jaeger, and W. W. Grabow, “Fluorescent monitoring of RNA assembly and processing using the split-spinach aptamer,” ACS synthetic biology, vol. 4, pp. 162–166, 2015.

[24] G. Pothoulakis, F. Ceroni, B. Reeve, and T. Ellis, “The spinach RNA aptamer as a characterization tool for synthetic biology,” ACS synthetic biology, vol. 3, pp. 182–187, 2014.

[25] R. Marshall and V. Noireaux, “Synthetic biology with an all e. coli txtl system: Quantitative characterization of regulatory elements and gene circuits,” in Synthetic Biology, ed: Springer, 2018, pp. 61–93.

[26] H. Niederholtmeyer, L. Xu, and S. J. Maerkl, “Real-time mRNA measurement during an in vitro transcription and translation reaction using binary probes,” ACS synthetic biology, vol. 2, pp. 411–417, 2013.

[27] T. Stögbauer, L. Windhager, R. Zimmer, and J. O. Rädler, “Experiment and mathematical modeling of gene expression dynamics in a cell-free system,” Integrative Biology, vol. 4, pp. 494–501, 2012.

[28] A. Adhikari, M. Vilkhovoy, S. Vadhin, H. E. Lim, and J. D. Varner, “Effective biophysical modeling of cell free transcription and translation processes,” Frontiers in bioengineering and biotechnology, vol. 8, 2020.

[29] R. Marshall and V. Noireaux, “Quantitative modeling of transcription and translation of an all-E. coli cell-free system,” Scientific reports, vol. 9, pp. 1–12, 2019.

[30] Y. Zhao, R. Yang, Z. Bousraou, and S. Wang, “Probing Human Osteogenic Differentiation Using Double-Stranded Locked Nucleic Acid Biosensors,” in 2021 IEEE 21st International Conference on Nanotechnology (NANO), 2021, pp. 273–276.

[31] R. Riahi, Z. Dean, T.-H. Wu, M. A. Teitell, P.-Y. Chiou, D. D. Zhang, et al., “Detection of mRNA in living cells by double-stranded locked nucleic acid probes,” Analyst, vol. 138, pp. 4777–4785, 2013.

[32] S. Wang, R. Riahi, N. Li, D. D. Zhang, and P. K. Wong, “Single cell nanobiosensors for dynamic gene expression profiling in native tissue microenvironments,” Advanced Materials, vol. 27, pp. 6034–6038, 2015.

[33] S. Wang, Y. Xiao, D. D. Zhang, and P. K. Wong, “A gapmer aptamer nanobiosensor for real-time monitoring of transcription and translation in single cells,” Biomaterials, vol. 156, pp. 56–64, 2018.

[34] R. L. Strack, W. Song, and S. R. Jaffrey, “Using Spinach-based sensors for fluorescence imaging of intracellular metabolites and proteins in living bacteria,” Nature protocols, vol. 9, pp. 146–155, 2014.

[35] W. Song, R. L. Strack, N. Svensen, and S. R. Jaffrey, “Plug-and-play fluorophores extend the spectral properties of Spinach,” Journal of the American Chemical Society, vol. 136, pp. 1198–1201, 2014.

[36] Z. Zhou, M. S. Hossain, and D. Liu, “Involvement of the long noncoding RNA H19 in osteogenic differentiation and bone regeneration,” Stem Cell Research & Therapy, vol. 12, pp. 1–9, 2021.

[37] V. Noireaux, R. Bar-Ziv, and A. Libchaber, “Principles of cell-free genetic circuit assembly,” Proceedings of the National Academy of Sciences, vol. 100, pp. 12672–12677, 2003.

[38] D. Garenne, M. C. Haines, E. F. Romantseva, P. Freemont, E. A. Strychalski, and V. Noireaux, “Cell-free gene expression,” Nature Reviews Methods Primers, vol. 1, pp. 1–18, 2021.

[39] H. Jia and P. Schwille, “Bottom-up synthetic biology: reconstitution in space and time,” Current opinion in biotechnology, vol. 60, pp. 179–187, 2019.

[40] N. Laohakunakorn, L. Grasemann, B. Lavickova, G. Michielin, A. Shahein, Z. Swank, et al., “Bottom-up construction of complex biomolecular systems with cell-free synthetic biology,” Frontiers in bioengineering and biotechnology, vol. 8, p. 213, 2020.

[41] A. P. Liu, “The rise of bottom-up synthetic biology and cell-free biology,” Physical biology, vol. 16, p. 040201, 2019.

[42] A. Groaz, H. Moghimianavval, F. Tavella, T. W. Giessen, A. G. Vecchiarelli, Q. Yang, et al., “Engineering spatiotemporal organization and dynamics in synthetic cells,” Wiley Interdisciplinary Reviews: Nanomedicine and Nanobiotechnology, vol. 13, p. e1685, 2021.

[43] H. Maeda, N. Fujita, and A. Ishihama, “Competition among seven Escherichia coli σ subunits: relative binding affinities to the core RNA polymerase,” Nucleic acids research, vol. 28, pp. 3497–3503, 2000.

[44] T. M. Gruber and C. A. Gross, “Multiple sigma subunits and the partitioning of bacterial transcription space,” Annual Reviews in Microbiology, vol. 57, pp. 441–466, 2003.

[45] G. Shin, “Molecular programming with a transcription and translation cell-free toolbox: from elementary gene circuits to phage synthesis,” 2012.

