## Supplemental Figures for "Experimental and Biophysical Modeling of Transcription and Translation Dynamics in Bacterial- and Mammalian-based Cell-Free Expression Systems"

**Table S1.** Terms of CFE systems

**Table S2**. Plasmid list

**Table S3**. Parameters for modeling

**Rate equations:** two-stage transcriptional activation cascade

**Figure S1.** Illustration of a simple biophysical model for mRNA and protein synthesis in CFE systems.

**Figure S2.** Calibration curve of broccoli aptamer.

**Figure S3.** Standard calibration curve of GFP.

**Figure S4.** Calibration curve of dsLNA probe.

**Figure S5.** Synthesize deGFP concentrations versus DNA (pTXTL-P70a-deGFP) concentrations in E coli-based CFE systems

**Figure S6.** Synthesized GFP concentrations versus DNA (pT7-CFE-GFP) concentrations in HeLa-based CFE systems.

**Figure S7.** Protein synthesis dynamics in HeLa-based CFE system with the duration of 10 hours.

**Table 1. Terms of CFE systems**

| Terms | Description | Source – references |
|---|---|---|
| P70a | Lambda phage promoter specific to E. coli σ70 | GenBank |
| P28a | Promotor of the E. coli tar chemotaxis gen to σ28 | Ref [1] |
| σ28 | E. coli sigma factor 28 | Ref [2] |
| Broccoli | Fluorescent RNA aptamer in present of DFHB1-T | Ref [3] |
| deGFP | eGFP truncated and modified in N-and C- terminus | Ref [4] |
| T7 | T7 bacteriophage RNA polymerase | Ref [5] |
| GADD34 | GADD34 truncated at N-terminus, translation enhancer | Ref [6] |

**Table 2. Plasmid list**

| Plasmids | Description |
|---|---|
| pTXTL-P70a-Broccoli | Single promotor sigma 70 to activate RNA apatamer |
| pTXTL-P70-deGFP | Single promotor sigma 70 to activate deGFP |
| pTXTL-P70a-S28 | Sigma 70 to activate the expression of sigma 28 |
| pTXTL-P28a-deGFP | Sigma 28 activate the expression of deGFP |
| pT7-CFE-GFP | T7 promotor to activate GFP |

### Table 3. Parameters for modeling

| Gene Circuits | Parameter | Value | Description |
|---|---|---|---|
| *E. coli*-based CFE<br>pTXTL-p70a-Broccoli | $\alpha_{tr}$ | 0.065 ( $s^{-1}$) | mRNA synthesis rate |
| | $\beta_{tr}$ | 6.6 nM $s^{-1}$ | Degradation rate |
| *E. coli*-based CFE<br>pTXTL-P70a-deGFP | $\alpha_{tr}$ | 0.065 ( $s^{-1}$) | mRNA synthesis rate |
| | $\beta_{tr}$ | 6.6 nM $s^{-1}$ | Degradation rate |
| | $\alpha_{tl}$ | 0.006 ( $s^{-1}$) | Translation rate |
| | $\beta_{tl}$ | 0 (nM/s) | Immature protein degradation rate |
| | $\alpha_{mp}$ | 0.000725 ($s^{-1}$) | Maturation rate |
| | $\beta_{mp}$ | 0 (nM/s) | Protein degradation rate |
| *E. coli*-based CFE<br>Two-stage cascade. | $\alpha_{tr_{70}}$ | 0.2 ($s^{-1}$) | S28 mRNA synthesis rate |
| | $\beta_{tr_{m28}}$ | 6.6 nM $s^{-1}$ | Degradation rate |
| | $\alpha_{tl}$ | 0.002 ($s^{-1}$) | Translation rate ( |
| | $\beta_{m_{28}}$ | 0 (nM/s) | S28 protein degradation rate |
| | $\alpha_{m_{28}}$ | 0.001 ($s^{-1}$) | S28 maturation rate |
| | $\alpha_{tr_{28}}$ | 0.04 ( $s^{-1}$) | deGFP mRNA synthesis rate |
| | $\beta_{tr_{meGFP}}$ | 0 (nM/s) | deGFP immature protein degradation rate |
| | $\alpha_{m_{eGFP}}$ | 0.001 ( $s^{-1}$) | deGFP maturation rate |
| | $\beta_{m_{eGFP}}$ | 0 (nM/s) | deGFP protein degradation rate |
| HeLa-based CFE<br>pT7-CFE-GFP | $\sigma_{tr}$ | 0.025 ($s^{-1}$) | mRNA synthesis rate |
| | $t_m$ | 1000 (s) | mRNA inactivation time |
| | $\sigma_{tl}$ | 0.001 ($s^{-1}$) | Translation rate |
| | $\beta_{mp}$ | 0 (nM/s) | Protein degradation rate |
| | $\sigma_{mp}$ | 0.003 ($s^{-1}$) | Maturation rate |

**Rate equations for two-stage transcriptional activation cascade**

$$\frac{dm_{28}}{dt} = \alpha_{tr_{70}}[DNA_{28}] - \beta_{tr_{m28}}[m_{28}]$$

$$\frac{dp_{28}}{dt} = \alpha_{tl}[m_{28}] - \beta_{m_{28}}[p_{28}]$$

$$\frac{dp_{28,m}}{dt} = \alpha_{m_{28}}[p_{28}]$$

$$\frac{dm_{egfp}}{dt} = \alpha_{tr_{28}}[p_{28,m}][DNA - eGFP] - \beta_{tr_{meGFP}}[m_{eGFP}]$$

$$\frac{dp_{eGFP}}{dt} = \alpha_{tl}[m_{eGFP}] - \beta_{m_{eGFP}}[p_{eGFP}]$$

$$\frac{dp_{eGFP,m}}{dt} = \alpha_{m_{eGFP}}[p_{eGFP}]$$

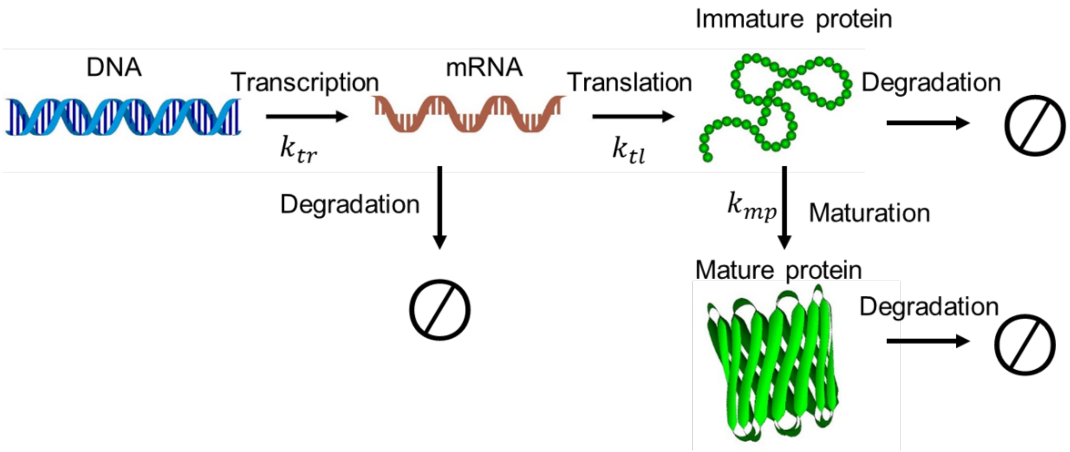


**Figure S1**. Illustration of a simple biophysical model for mRNA and protein synthesis in CFE systems. The model describes the gene expression dynamics including transcription, translation, and protein maturation.

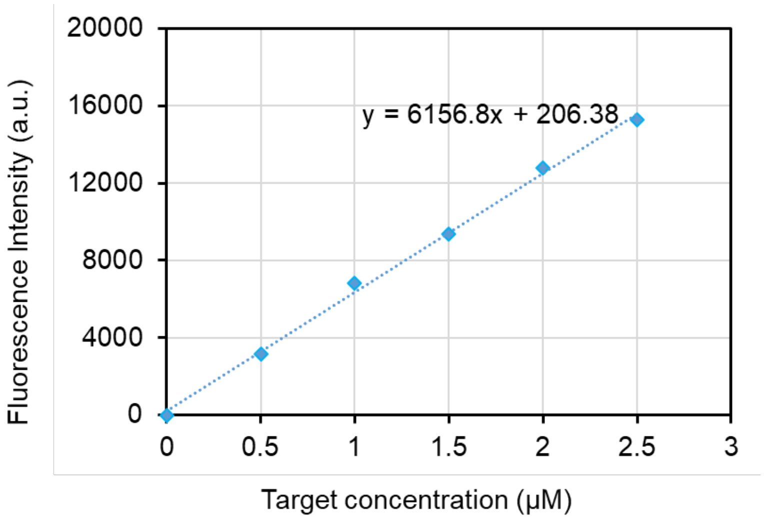


**Figure S2**. Calibration curve of broccoli aptamer. The curve was generated by measuring fluorescence intensity of DNA oligos (broccoli oligo and its target oligos) at varying concentrations in a microplate reader. The excitation and emission wavelengths are 470 nm and 505 nm, respectively. Experiments were performed three times independently with triplicates. Data are presented as mean ± s.e.m. (n = 3).

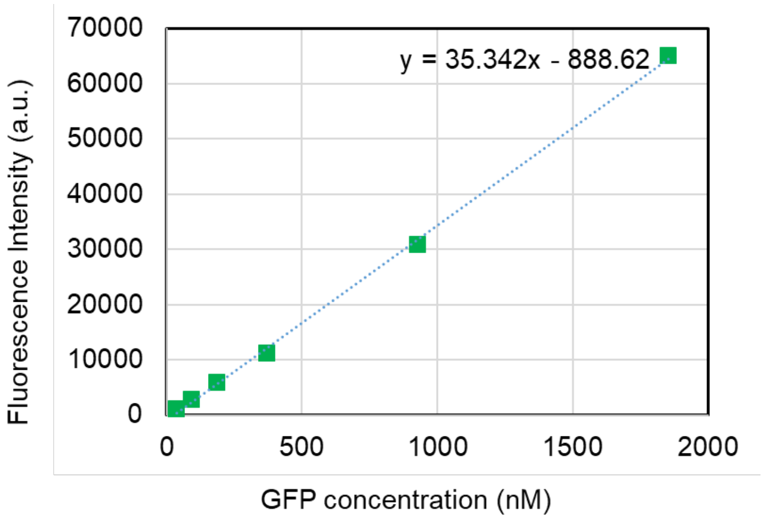


**Figure S3**. Standard calibration curve of GFP. The standard curve was generated by measuring fluorescence intensity of GFP at varying concentrations in a microplate reader. The excitation and emission wavelengths are 485 nm and 525 nm, respectively. Experiments were performed three times independently with triplicates. Data are presented as mean ± s.e.m. (n = 3).

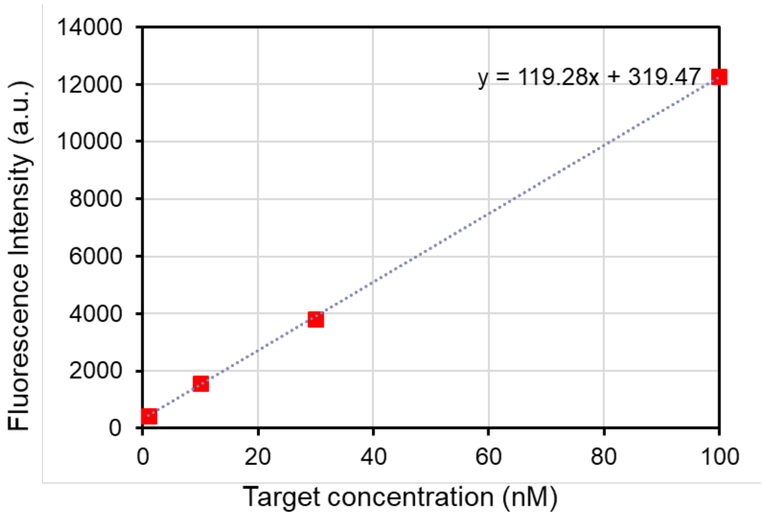


**Figure S4**. Calibration curve of dsLNA probe. The calibration curve was generated by measuring fluorescence intensity of dsLNA probe with different concentrations of target (DNA oligos). The excitation and emission wavelengths are 485 nm and 525 nm, respectively. Experiments were performed three times independently with triplicates. Data are presented as mean ± s.e.m. (n = 3).

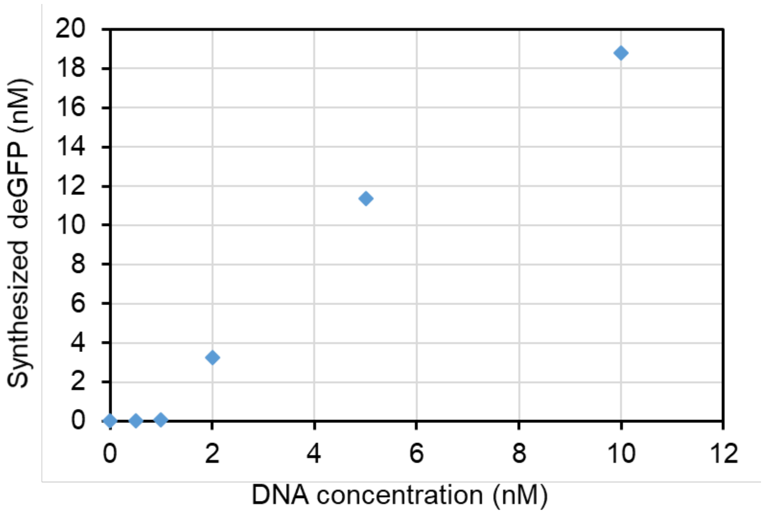


**Figure S5**. Synthesize deGFP concentrations versus DNA (pTXTL-P70a-deGFP) concentrations in *E coli*-based CFE systems. Data were obtained at the end point of 3 hr incubation.

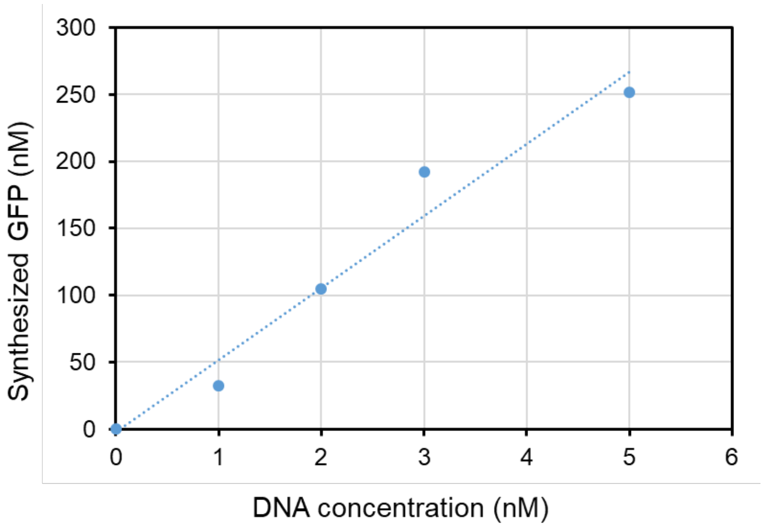


**Figure S6**. Synthesized GFP concentrations versus DNA (pT7-CFE-GFP) concentrations in HeLa-based CFE systems. Data were obtained at the end point of 4 hr incubation. Data represents at least three independent repeats with triplets.

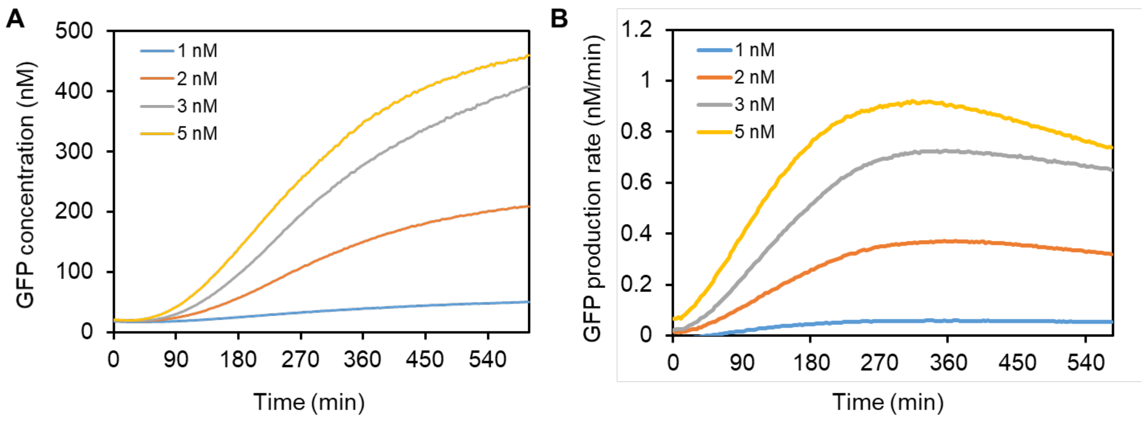


**Figure S7**. Protein synthesis dynamics in HeLa-based CFE system with the duration of 10 hours. **(A)** Synthesized GFP concentrations with plasmid DNA concentrations of 1 nM, 2 nM, 3 nM, and 5 nM over the period of 10 hours. **(B)** GFP protein rate with plasmid DNA concentrations of 1 nM, 2 nM, 3 nM, and 5 nM over the period of 10 hours.

### References


[1] D. N. Arnosti and M. J. Chamberlin, "Secondary sigma factor controls transcription of flagellar and chemotaxis genes in Escherichia coli," *Proceedings of the National Academy of Sciences,* vol. 86, pp. 830-834, 1989.

[2] J. Shin and V. Noireaux, "An E. coli cell-free expression toolbox: application to synthetic gene circuits and artificial cells," *ACS synthetic biology,* vol. 1, pp. 29-41, 2012.

[3] G. S. Filonov, J. D. Moon, N. Svensen, and S. R. Jaffrey, "Broccoli: rapid selection of an RNA mimic of green fluorescent protein by fluorescence-based selection and directed evolution," *Journal of the American Chemical Society,* vol. 136, pp. 16299-16308, 2014.

[4] J. Garamella, R. Marshall, M. Rustad, and V. Noireaux, "The all E. coli TX-TL toolbox 2.0: a platform for cell-free synthetic biology," *ACS synthetic biology,* vol. 5, pp. 344-355, 2016.

[5] S. Wang, S. Majumder, N. J. Emery, and A. P. Liu, "Simultaneous monitoring of transcription and translation in mammalian cell-free expression in bulk and in cell-sized droplets," *Synthetic Biology,* vol. 3, p. ysy005, 2018.

[6] S. Mikami, T. Kobayashi, K. Machida, M. Masutani, S. Yokoyama, and H. Imataka, "N-terminally truncated GADD34 proteins are convenient translation enhancers in a human cell-derived in vitro protein synthesis system," *Biotechnology letters,* vol. 32, pp. 897-902, 2010.
